# Brain transcriptomic changes in Japanese quail reveal roles for neurotensin and urocortin 3 in avian parental care

**DOI:** 10.1101/2020.11.29.401794

**Authors:** Patricia C. Lopes, Robert de Bruijn

**Author notes:** Corresponding author Patricia C. Lopes.

## Abstract

For many species, parental care critically affects offspring survival. But what drives animals to display parental behaviours towards young? In mammals, pregnancy-induced physiological transformations seem key in preparing the neural circuits that lead towards attraction (and reduced-aggression) to young. Beyond mammalian maternal behaviour, knowledge of the neural mechanisms that underlie parental care is severely lacking. We took advantage of a domesticated bird species, the Japanese quail, for which parental behaviour towards chicks can be induced through a sensitization procedure, a process that is not effective in all animals. We used the variation in parental responses to study neural transcriptomic changes associated with the sensitization procedure itself and with the outcome of the procedure (i.e., presence of parental behaviours). Out of the brain regions studied, we found that most differences in gene expression were located in the hypothalamus. Two genes identified are of particular interest, as no role in avian parental care was known for those genes. One is neurotensin, previously only demonstrated to be causally associated with maternal care in mammals. The other one is urocortin 3, causally demonstrated to affect young-directed neglect and aggression in mammals. Our work opens new avenues of research into understanding the neural basis of parental care in non-placental species.

## 1. Introduction

Most of what is known regarding the mechanistic basis of infant-directed parental care comes from studies in mammals [1]. In mammals, pregnancy-induced physiological transformations seem key in preparing the neural circuits that lead towards attraction (and reduced-aggression) to an infant [1]. However, not all parental care is explained by pregnancy-induced changes. For example, there is also paternal care, alloparenting (parental care towards non-descendant young including adoption) and care by non-placental animals [2]. Knowledge of what neurotransmitters/neuromodulators are involved in these situations is lacking. While these forms of care are widespread in human societies [3] and in non-placental vertebrates [4], and play critical roles in offspring development [5,6], their neural underpinnings are severely understudied. Studying species that show parental care in the absence of pregnancy can therefore help elucidate the neurochemical make-up of these diverse forms of parental care.

While parental care is rare in many vertebrate taxa, it is essential for survival in the majority of avian species [7]. Although biparental care is the most frequent modality observed, birds as a group show an incredible diversity in forms of parental care, from brood parasitism, where care is lost, to cooperative breeding, where non-reproductive individuals provide care to offspring that are not their own [7]. In terms of neuroendocrine mechanisms associated with incubation and caring for young, the majority of attention has been paid to a single pituitary hormone: prolactin [8,9]. Coupled with an increase in prolactin, a decline in gonadal hormones seems to facilitate the transition between courtship and nest building into incubation behaviour [10]. The neural underpinnings of young-directed parental care in birds are, however, severely understudied.

Using a sensitization procedure developed by [11], a recent experiment by our group confirmed that it is possible to induce parental behaviours in Japanese quail through a single overnight exposure to chicks [12]. After the sensitization procedure, both sexes display chick brooding behaviour and reduced aggression towards chicks (pecking behaviour). This procedure therefore allows us to study young-directed care at its onset, in both sexes, in the absence of pregnancy-induced effects, copulation or pharmacological hormonal priming. What is also interesting is that, while the change in behaviour after sensitization is drastic, it is not observed in all animals, i.e., some animals fail to respond to the sensitization. Conversely, a few animals not exposed to the sensitization spontaneously show some parental care when paired with chicks.

In the current study, we take advantage of this variation in parental care to understand, from a neural molecular perspective, a) what modifications occur due to the sensitization treatment, b) what genes correlate with the duration of brooding behaviour and with chick-directed aggression, and c) whether the same mechanisms are involved in care that is shown after sensitization treatment relative to care that is shown spontaneously (i.e., in the absence of sensitization treatment). We focused our transcriptomic analysis on the hypothalamus, the bed nucleus of the stria terminalis and the nucleus taeniae. We targeted these brain regions because they contain nuclei important for parental care, affiliative interactions, and agonistic interactions in birds and other vertebrates [2,4,13–16]. Our experiments highlight the importance of using intra-species variation in parental responses to elucidate neural mechanisms of parental behaviour and to discover new molecular pathways involved in avian parental care, several of which are likely conserved across other vertebrate taxa.

## 2. Materials and methods

### (a) Parental care induction procedure and behaviour quantification

Japanese quail (*Coturnix japonica*) were raised in the lab from eggs obtained from AA Lab Eggs, Inc. (Westminster, CA) following the procedures described in [12]. Adult birds used in this experiment were kept in an 8L:16D light cycle, at a temperature range of 20-24 °C and maintained in groups of 4-5 same sex animals in cages (100 × 40 × 50 cm). We chose a short photoperiod for this experiment for two reasons: 1) male and female Japanese quail kept at short photoperiods are reproductively quiescent [17], which reduces the influence of gonadal hormones from the study; and 2) quail sensitization treatments are more effective at increasing brooding duration in both sexes under short photoperiods [11,16]. The reason for more effective sensitization when gonadal hormones are low may be linked to the natural decline in gonadal hormones undergone by birds transitioning between courtship and nest building to a parental state (onset of incubation) (reviewed in [10]). Two days prior to carrying out the experiments, a replica of the wooden box (18 × 18 × 18 cm) used in the parental care sensitization procedure was added to those cages. The day prior to the procedure, animals were separated individually into cages identical to the group cages, which also contained a wooden box. The wooden box was always open during these days, allowing the birds to inspect it and go in and out. Food and water were provided *ad libitum* during the entire experiment.

The day of the parental care sensitization procedure, each adult bird was locked inside the wooden box in their individual cage starting at one hour before lights were off. Under the parental induction treatment, two chicks (1-3 days old) were added to this box just before lights off. Under the control treatment, no chicks were added to the box. Animals were left undisturbed overnight. The morning after, the wooden box was opened, the overnight chicks were removed, and two new chicks (1-3 days old) were added to the cage. This swap of chicks ensured that birds in both treatments (controls and sensitized) were being tested for their behaviours towards novel chicks (in case familiarity influenced the outcome). The novel chicks were allowed to stay in the cage for 20 min, during which videos were continuously recorded (Axis M1065L network camera by Axis Communications). At the end of the 20 min, adults were euthanized by isoflurane inhalation and decapitation. The brain was removed from the skull, flash frozen and then stored at −80 °C until further processing. Chicks were returned to their cages. In total, 50 adults (ages 90-110 days old) were subjected to the experimental procedures, of which 11 females and 13 males were randomly assigned to the sensitization treatment and 19 females and 7 males to the control treatment. Two control females were removed from the study due to methodological failures during the experiments, bringing the final number of control females to 17. In total, 77 chicks were used. Behaviours towards chicks within the 20 min period of observation were scored by observers blind to the treatment and analysed as described in [12]. The behavioural results described here were re-plotted from [12] and were not re-analysed.

### (b) Brain dissection

Brains were coronally sectioned on a Leica CM1860UV cryostat, at −18 °C. We used separate surgical micropunches (EMS Rapid Core Instruments) to punch out the three brain regions of interest from 100□μm slices, spaced apart by three 30□μm slices collected onto microscope slides (Fisherbrand, item 12–550-15) for future use. The brain regions collected were the entire hypothalamus, the nucleus taeniae (Tn) and the bed nucleus of the stria terminalis (BnST). These regions were identified based on use of both the quail and the chicken brain atlases [18,19]. The diameter of the micropunchers used was 4 mm for the hypothalamus and 3 mm for the other regions. The hypothalamus was collected from the start of the bifurcation the tractus septomesencephalicus until the start of the posterior commissure. We started collecting the BnST from the point when the anterior commissure was visible, using the lateral ventricle as a guide for positioning the micropuncher, and ended after collecting 5 punches, i.e., over 1 cm coronally (as each punch is separated from the next by about 190 mm). Collection of the Tn started at the point where the occipitomesencephalic tract splits into two and ended after 5 punches (again equivalent to 1 cm coronally). Punches from each of these brain regions were placed in individual tubes containing 2□mm size beads (ZR BashingBeads Lysis Tubes, Zymo Research, item S6003-50) and 1 mL of QIAzol lysis reagent (Qiagen, item #79306). Tissue homogenization was done by agitating the tubes for 20□s at a 7□m□s^−□1^ speed (Beadbug 6 homogenizer, Benchmark Scientific), followed by a 5□min rest period. The homogenate was then transferred into a new tube and preserved at − □80□°C until RNA isolation.

### (c) RNA isolation, library preparation and sequencing

Total RNA was extracted from the aqueous layer formed after chloroform precipitation, using the RNA Clean & Concentrator Kit-5 (Zymo Research, item # R1013) following manufacturer’s instructions and including the DNase I in-column treatment step. These samples were then shipped on dry ice to Novogene Corporation Inc. (Chula Vista, CA, USA), where RNA quantity, purity and integrity were assessed on an Agilent 2100 Bioanalyzer (Agilent Technologies, Santa Clara, CA), cDNA libraries were generated using NEBNext® UltraTM RNA Library Prep Kit for Illumina® (NEB, USA) and cDNA fragments (150 ∼ 200 bp in length) were then purified using the AMPure XP System (Beckman Coulter, Beverly, USA). Paired-end sequencing of libraries (PE150; Illumina Novaseq 6000) was performed according to standard protocols. An average of 52 million paired-end raw reads were obtained for each sample.

### (d) Mapping and differential gene expression analysis

An average of 85.3% of clean (post adapter removal and quality filtering) reads were mapped to the Japanese quail reference genome (Coturnix_japonica_2.0, INSDC Assembly Mar 2016, downloaded from Ensembl), representing an average of 43.7 million mapped reads per sample (Table S1 contains information on mapping statistics per sample). Mapping was done using HISAT2 [20], and HTSeq was used to count the number of mapped reads to each gene [21]. Differential gene expression analysis for the effect of treatment was performed using the DESeq2 R package [22,23]. To determine whether any genes showed significant correlations to either brooding duration or aggressive events, we used edgeR TMM-normalized counts (edgeR package [24]) with the limma package [25], with voom transformation [26]. While the DESeq2 package can also accept continuous covariates in the model, the list of significant genes detected using the limma+voom approach overlaps with the one produced by DESeq2 but excludes genes for which the regression line does not properly fit the points when plotted. To control for the false discovery rate due to multiple testing, *p*-values were adjusted using the Benjamini-Hochberg procedure. Genes were considered as statistically differentially expressed when adjusted *p*-values were□<□0.05.

To examine the effect of treatment on gene expression, for each brain region, we first tested whether the variable sex interacted with the variable treatment. In case no differentially expressed genes (DEGs) were detected for the interaction term at p_adj_ < 0.05, the variable sex was excluded from the model and treatment was tested alone. To examine covariation between gene expression levels and our two continuous variables (brooding duration or aggression), for each brain region, we first tested for an interaction of treatment or sex with each continuous covariate. In case no DEGs were detected for the interaction term at p_adj_ < 0.05, the variable treatment was excluded from the model and the continuous covariates were tested alone. Prior to testing, the continuous covariates were scaled and centered, but figures show brooding duration in seconds for simplicity of interpretation. Volcano plots for the results of treatment effects were produced using log2 transformed DESeq2 normalized counts. A subset of representative genes obtained from the limma-voom analysis were represented by plotting log2 voom transformed TMM normalized counts, and a regression line with a slope of the logFC and the coefficient for the intercept for each specific gene.

### (e) Gene ontology (GO) enrichment analysis and KEGG pathway enrichment analysis of differentially expressed genes

Functional enrichment analysis was done only for the DEGs found in the hypothalamus, as this region contained the most DEGs both for the effect of treatment and for the covariation with parental duration. The stringApp (v. 1.5.4) [27] within Cytoscape software (v. 3.8.2) [28] was used to find enriched GO terms and enriched KEGG Pathways. Up- and down-regulated DEGs were analysed separately. *Mus musculus* was used as the background species, and the significance threshold for the false discovery rate (FDR) was 0.05. We used the same app and settings to visualize possible protein-protein interactions (PPI) among the protein products of these DEGs, but this time we analysed all genes (up and down-regulated) at once. This analysis produces a PPI enrichment p-value, which, when smaller or equal than 0.05, indicates that the proteins being tested have more interactions among themselves than would be expected for a random set of proteins of similar number, drawn from the genome.

## 3. Results

### (a) Effect of sensitization treatment

#### Behavioural response

In response to spending an overnight enclosed in a box with two chicks (sensitization treatment), the average duration of time spent brooding novel chicks was significantly increased and not different by sex (Fig. 1a, [12]). Out of animals in the sensitization treatment, 3 out of 11 females (27.3 %) and 3 out of 13 males (23.1 %) showed no brooding behaviour. Out of control animals, 6 out of 16 females (37.5 % of females) and 3 out of 8 males (37.5 % of males) showed at least a few seconds of brooding behaviour. Aggression, quantified as chick-directed pecking, was overall low and while it tended to be lower in animals in the sensitization treatment, this difference was not significant (Fig. 1b, [12]).

**Figure 1.**
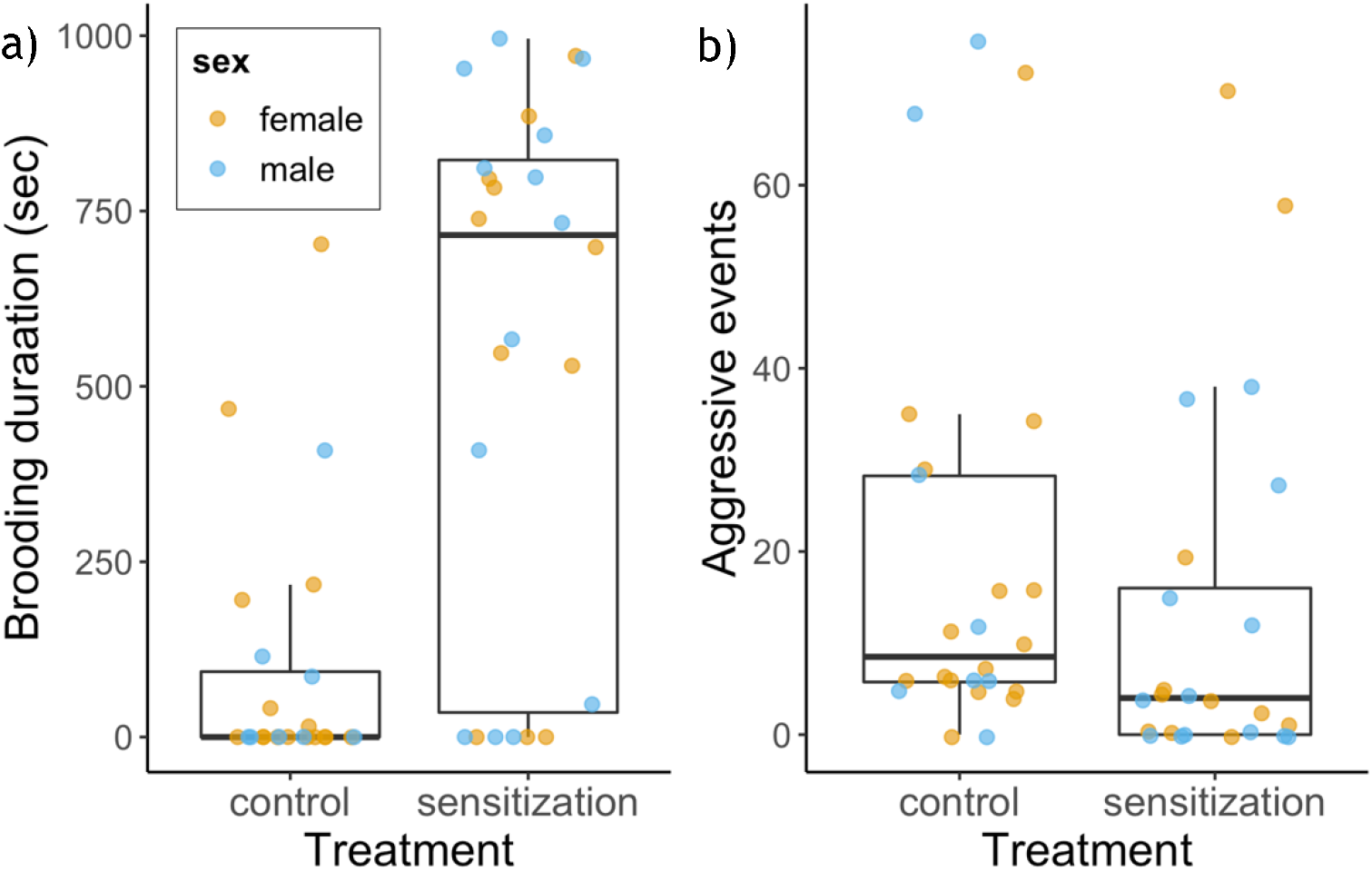
Brooding duration (a) and aggressive events (chick-directed pecking) (b) in animals that underwent sensitization treatment or not (control). Dots represented individual animals, coloured by sex.

#### Transcriptomic response

Sex did not interact with treatment in affect gene expression in any of the brain regions. Therefore, the effect of treatment was tested in both sexes together. Very few genes were differentially expressed in the brains of animals that underwent the sensitization treatment relative to controls during interaction with chicks (Fig. 2, Table S2). In the hypothalamus, only two genes were up-regulated in the sensitization treatment relative to control: neurotensin (NTS) and urocortin 3 (UCN3). In the bed nucleus of the stria terminalis (BnST), only one gene was up-regulated: adipocyte plasma membrane associated protein (APMAP). This same gene was the only one that approached significance (padj = 0.07229) in the Nucleus Taeniae (Tn). Out of the six genes down-regulated in the hypothalamus of animals that underwent the sensitization treatment, three overlapped with down-regulated genes in the BnST: Fos proto-oncogene, AP-1 transcription factor subunit (FOS), early growth response 4 (EGR4) and BTG anti-proliferation factor 2 (BTG2). In addition to these three, another five genes were down-regulated in the BnST: Kruppel like factor 2 (KLF2), neuronal PAS domain protein 4 (NPAS4), RRAD, Ras related glycolysis inhibitor and calcium channel regulator (RRAD), salt inducible kinase 1 (SIK1) and Jun proto-oncogene, AP-1 transcription factor subunit (JUN).

**Figure 2.**
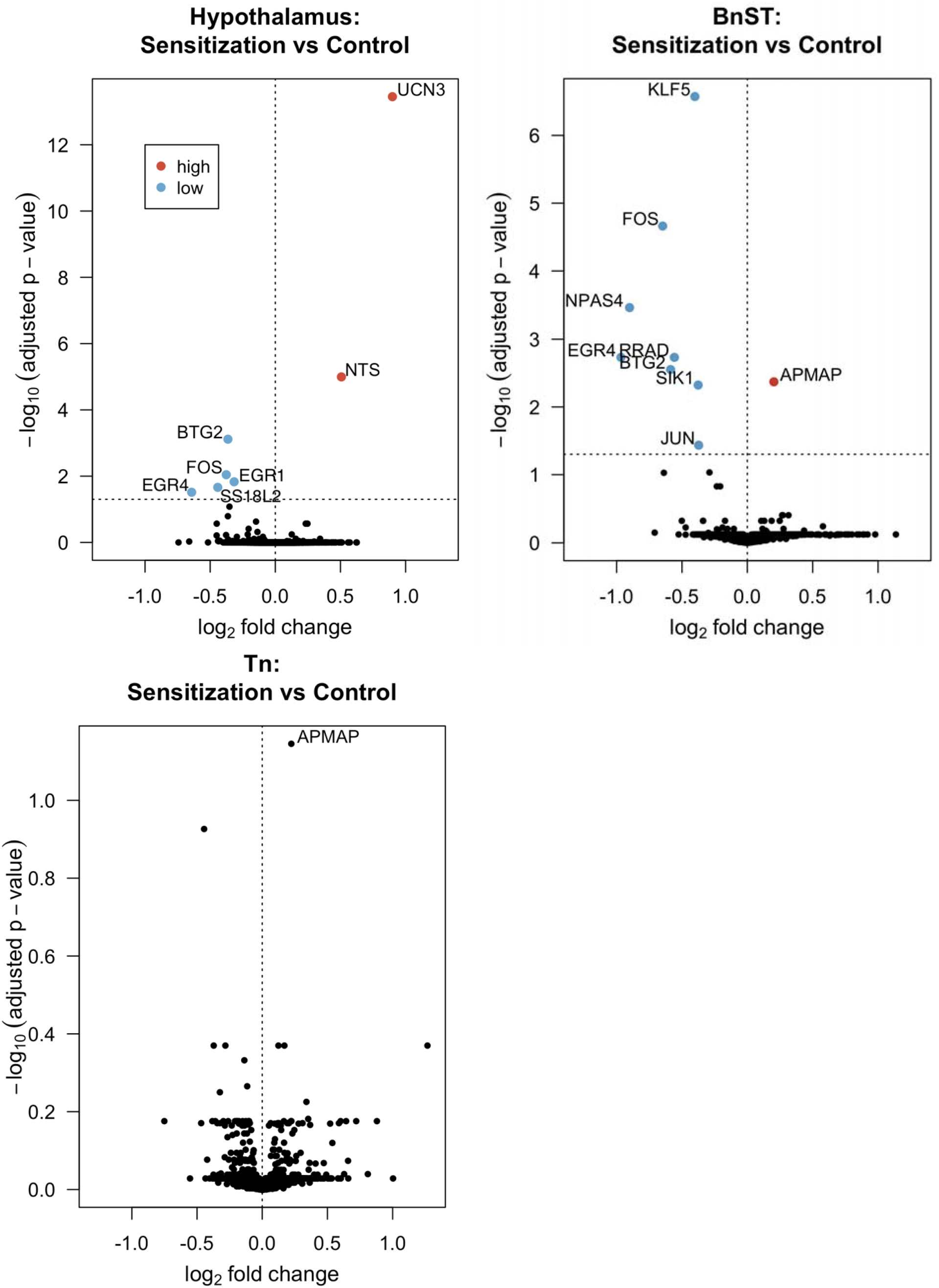
Volcano plots representing the log_2_ fold change against the corresponding – log_10_ p_adj_-value of all genes for each of the three brain regions studied. Differentially expressed genes (p_adj_ < 0.05) are shown in red if they are up-regulated in the sensitization treated animals relative to control and in blue if they are down-regulated. Other genes are shown in black. The horizontal dotted line indicates the p_adj_-value cut-off, while the vertical dotted line marks the point for zero fold change differences. One down-regulated gene without annotation is not show for hypothalamus (see Table S2 for complete list).

#### (b) Genes and pathways associated with the duration of brooding behaviour

The only brain region where genes showed a significant correlation with the duration of brooding behaviour was the hypothalamus. Twelve genes in this region were significantly correlated with brooding behaviour (Table S3). With the exception of transducer of ERBB2, 1 (TOB1), a down-regulated gene, all other genes overlap with DEGs detected in the hypothalamus or BnST. Furthermore, DEGs that were up-regulated in animals in the sensitization treatment relative to control increase in expression as brooding duration increases (UCN3, Fig. 3) and down-regulated genes decrease in expression as brooding duration increases (e.g., FOS, Fig. 3; other down-regulated genes show the same pattern). No genes were correlated with aggressive events in any of the brain regions studied. No interaction between brooding duration or aggressive events and sex was found. We also found no interaction between treatment and parental duration or treatment and aggressive events, indicating treatment did not affect the expression of genes related to either of these traits. In other words, genes that covary with parental duration are the same between birds in the sensitization treatment and birds in the control treatment and, therefore, the molecular mechanisms behind spontaneous care and care after undergoing the sensitization treatment are similar.

**Figure 3.**
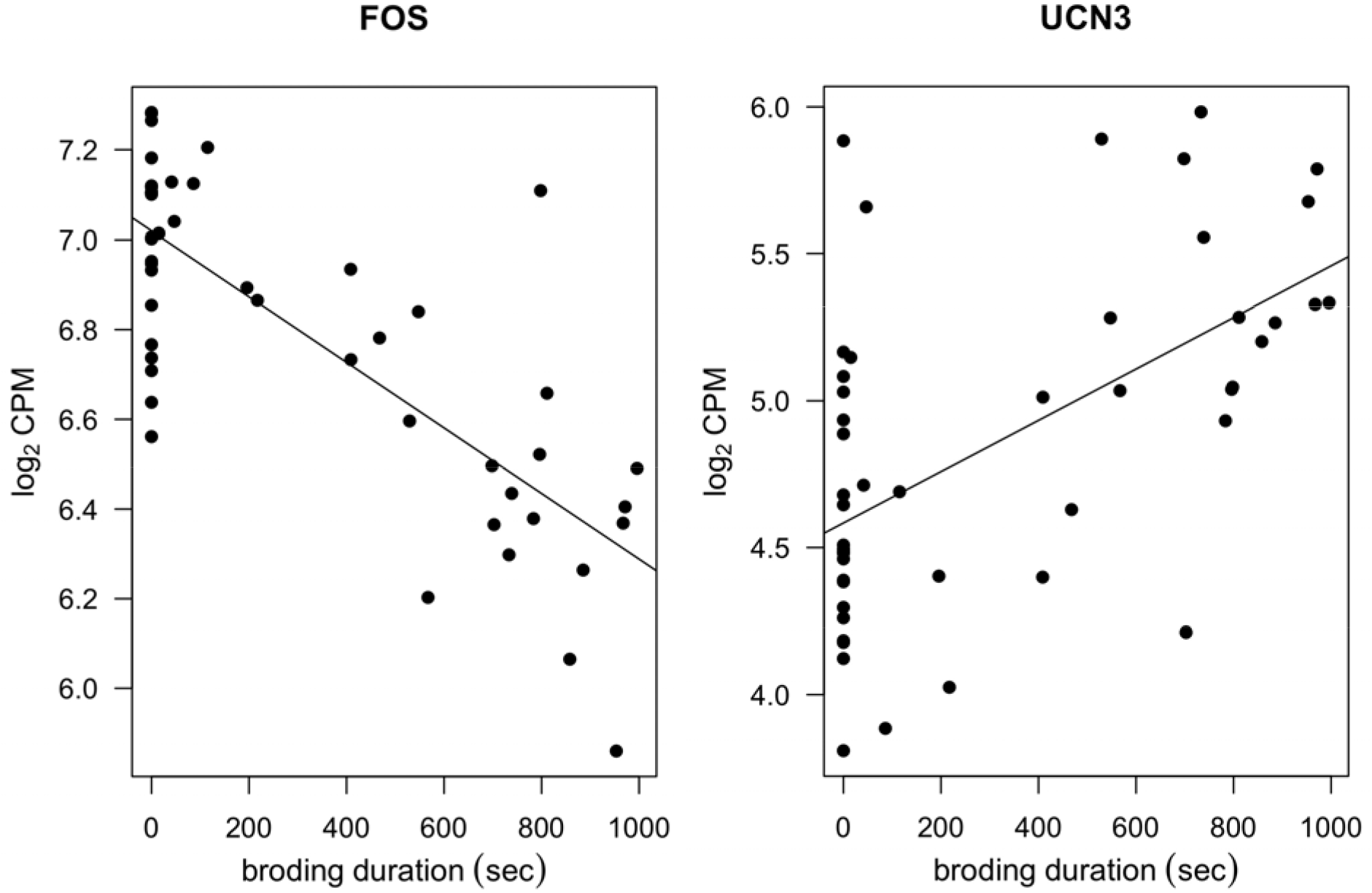
Regression lines representing the relationship between brooding duration (sex) and FOS expression (left panel) or UCN3 expression (right panel). Statistical details for all genes that covary with brooding duration can be found in Table S3.

To gain further insight into what pathways might be involved in brooding behaviour, we used GO enrichment analysis and KEGG pathway enrichment analysis (summary of results in Table I; full results in Tables S4 and S5). The GO enrichment analysis of DEGs that negatively covary with brooding behaviour revealed enrichment in several GO categories related with transcriptional regulation (most significant terms across all three GO categories), response to hormones (examples of terms with non-redundant gene combinations: response to corticosterone, response to steroid hormone, response to progesterone, cellular response to hormone steroid stimulus, response to peptide hormone) and with learning or memory (Table S4). In terms of genes that were either up-regulated by the sensitization treatment (UCN3 and NTS) or that positively covaried with brooding duration (UCN3), enriched GO terms were related to neuropeptide hormone activity (MF category) and with axon terminus and extracellular region (CC category).

**Table I.**
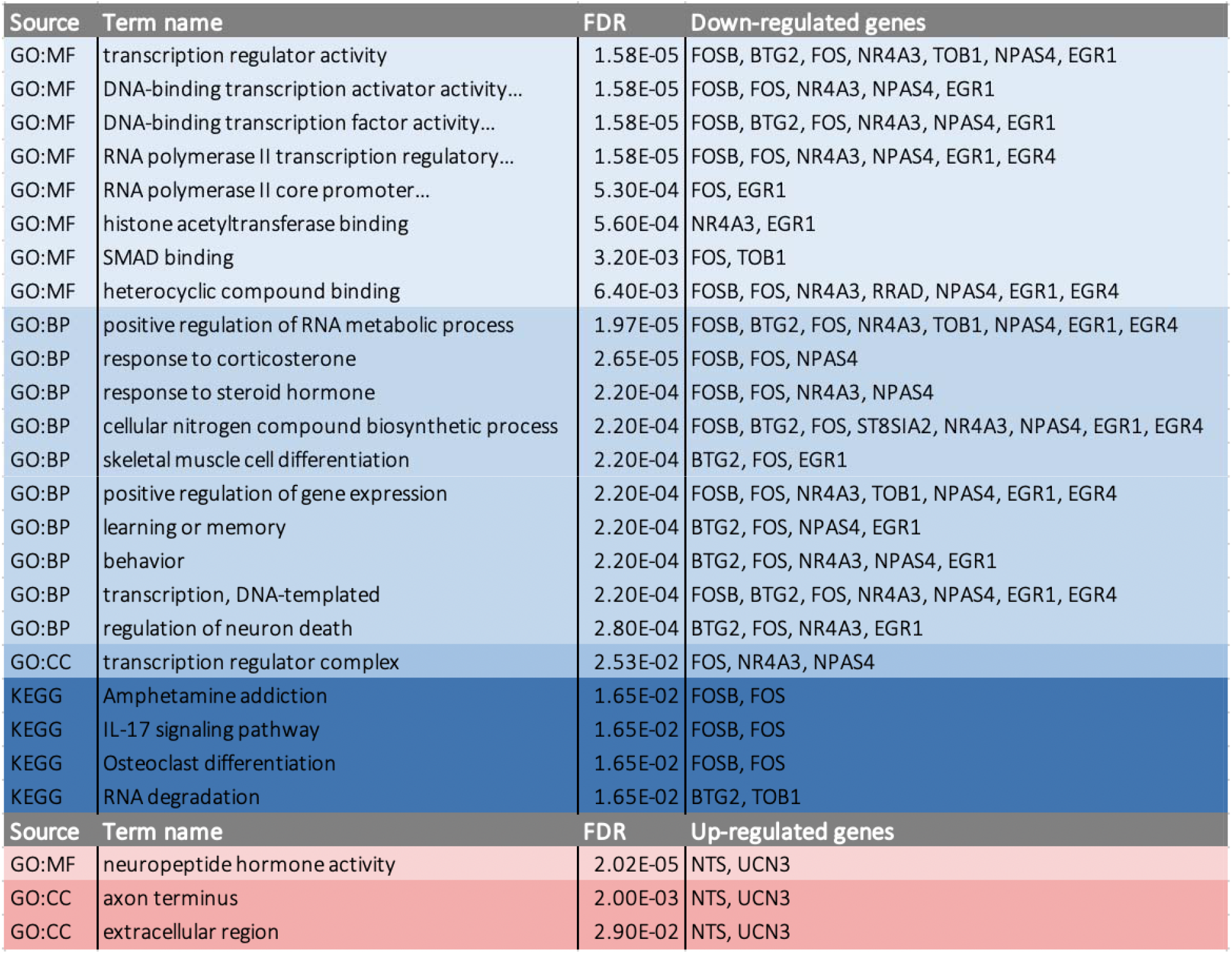
Significantly enriched GO terms and KEGG pathways associated with down- or up-regulated genes in the hypothalamus. Only top 10 terms per category are shown.For MF and BP, filtering of terms for redundancy of enriched genes was done (maximum Jaccard similarity of 1 allowed). Several term names are cropped for space (see Tables S4 and S5 for full list of unfiltered terms and complete term names). MF = Molecular function; BP = Biological Process; CC = Cellular Component.

In order to understand how the protein products of the DEGs may interact, if at all, we produced a protein-protein interaction network using the StringApp plugin within Cytoscape. The network produced showed that the majority of down-regulated genes interacted with one another and that NTS may interact with FOS (Fig. 4). The PPI enrichment p-value for this set of proteins is 1 × 10^−16^, indicating a larger degree of interaction among these proteins than would be expected by chance.

**Figure 4.**
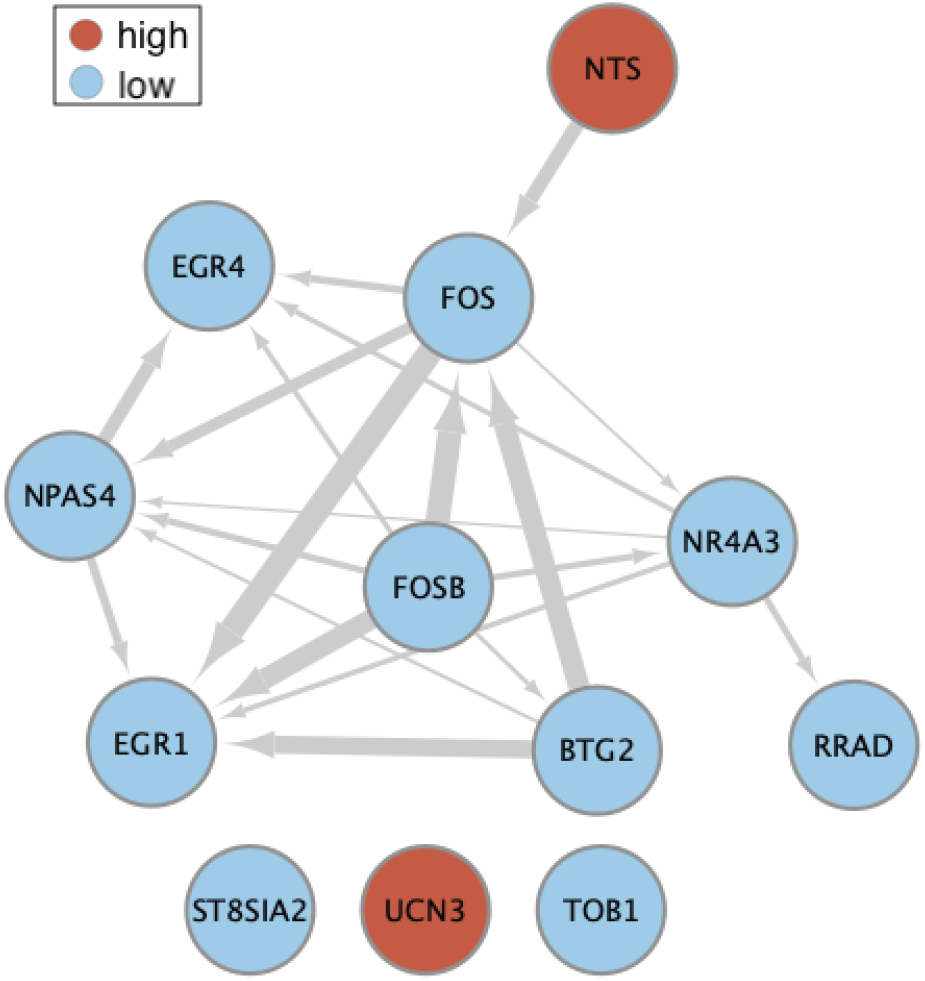
Protein-protein interaction network produced with STRING. Nodes with blue colours have lower expression in animals showing increased parental behaviours. Nodes with red colours have higher expression in animals in the sensitization treatment (both NTS and UCN3) or in animals showing increased parental behaviours (UCN3 only). The width of the edges (arrows) corresponds to the stringdb score, which measures how likely the interactions between two given proteins are to be true given the available evidence. In our network, we only show edges with medium (0.4) or better confidence.

## 4 Discussion

The neural mechanisms mediating parental care remain poorly understood. We took advantage of a domesticated species, the Japanese quail, for which parental behaviour towards chicks does not tend to occur spontaneously in captivity. Parental behaviour in these animals can be induced through a sensitization procedure, but the process is not 100 % effective. Using the variation in parental responses, we asked what neural molecular mechanisms: a) are altered upon sensitization treatment, b) correlate with the amount of parental response shown, and c) differ between care shown after the sensitization procedure relative to care shown in the absence of this procedure (spontaneous care). Out of the three brain regions studied, the only one where gene expression patterns correlated with the duration of brooding behaviour was the hypothalamus. Our unbiased transcriptomic approach identified two genes in this region, neurotensin and urocortin 3, that were previously only demonstrated to be causally associated with young-directed care and young-directed aggression in mammals. Transcriptomic studies focused on animals that fail to show parental care under care-promoting circumstances, or studies on those animals that show care unprompted provide great opportunities to understand the neural basis of variation in parental care.

By comparing the neural transcriptomic responses of adult quail that had undergone a sensitization procedure to those that did not undergo this process, we were able to identify genes that may be involved in activating avian parental care. One of those genes, neurotensin, was expressed at higher levels in the hypothalamus of both females and males after sensitization treatment. This finding is significant given that neurotensin levels in the hypothalamus of postpartum female mice were also observed to be elevated relative to that of virgin mice [29]. Neurotensin gene expression was also recently shown to be elevated in the preoptic area of males of one species of poison frog (*Dendrobates tinctorius*) when performing parental care relative to males not performing care [30]. Furthermore, intracerebroventricular injections of neurotensin significantly decreased maternal aggression in female mice [31], indicating the neurotensin participates in at least one component of maternal responses in rodents. Previously, the only reports of behavioural effects of neurotensin on birds suggested a possible relationship between neurotensin expression in the medial preoptic nucleus (POM) and non-vocal courtship behaviour and agonistic behaviour in male European starlings (*Sturnus vulgaris*) [32]. Interestingly, that study also reported that male starlings that had acquired a nest box also had higher neurotensin expression in the POM relative to males that did not acquire boxes and the measures of non-vocal courtship behaviour were mostly related to nest building (e.g., such as number of times entering or landing on the nest and gathering nesting material). European starlings show biparental care and the increase in neurotensin in a hypothalamic nucleus in that study could have been related to nest-building activity, which, for many species, is a component of parental behaviour.

Out of the hormones associated with avian parental care to date, prolactin is by far the one that has received the most attention [9]. In precocial avian species, as is the case with Japanese quail, prolactin tends to be elevated in females during the incubation period and to decline when chicks hatch [8]. Exposure to prolactin maybe therefore be needed to promote young-directed care. Prolactin release is known to be under the control of dopamine, which can have both inhibitory and stimulatory effects depending on the dopamine receptor subtype it acts on [9]. Neurotensin is capable of enhancing dopamine release [33] and could, in this way, impact parental behaviour by playing a role in controlling prolactin secretion. Neurotensin could also affect care through its involvement in reward processing [34].

Galanin is a neuropeptide that has emerged as an important regulator of parental behaviour [35]. While galanin was not detected in our study, it is interesting to note that neurotensin neurons in the medial preoptic area of the hypothalamus overlap with galanin neurons in this region [36]. This overlap may be suggestive of overlapping functions for the neurotensin neurons. It is also important to highlight that whereas neurotensin was elevated in birds exposed to the sensitization treatment, it did not covary with the extent of parental behaviour shown. A recent study that examined neurogenomic responses in paternal sticklebacks (*Gasterosteus aculeatus*) across several stages of care found that galanin, for example, was elevated in early stages (prior to hatching) but no different from controls once all eggs had hatched [37], highlighting that the role of different neuromodulators changes with the stage of care. Galanin may therefore be more important at a different stage of parental care in birds. Also, further studies will therefore be needed to understand whether neurotensin participates only in switching on parental behaviours (as is indicated by our study) or whether it is needed for the maintenance of those behaviours.

The nonapeptides oxytocin and vasopressin have been implicated in the regulation of social bonding and maternal care in mammals [1]. In birds, two studies (one in female turkeys, *Meleagris gallopavo* [38] and one in female Native Thai chickens, *Gallus domesticus* [39]) focused specifically on the transitions from egg-laying to incubation and from incubation to brooding behaviour, showed similar patterns in terms of number of neurons producing mesotocin (the avian homologue to oxytocin). Both studies found that the number of mesotocin-immunoreactive neurons was higher in specific hypothalamic nuclei of incubating than of non-incubating laying hens. While in the first chicken study showed no major differences in mesotocin-immunoreactive neurons between incubating and chick-brooding hens, a subsequent study on the same species found that hens that had been incubating eggs naturally and had their eggs swapped by chicks, had higher numbers of neurons showing mesotocin-immunoreactivity than hens that continued incubating eggs and received no chicks [40]. Combined, these studies suggest that mesotocin is important for transitions between parental care stages in some bird species. As we did not detect differences in mesotocin in our study, it may be that animals need to be actively laying or incubating eggs to express these changes in mesotocin. It will be interesting for future studies to explore these relationships further to understand the precise roles played by mesotocin in avian parental care behaviours.

The only other gene that was upregulated in the hypothalamus following the sensitization procedure was urocortin 3. Urocortin 3 was also the only gene in the hypothalamus to be positively correlated with brooding duration. In rodents, central administration or overexpression of urocortin 3 increases stress-induced anxiety and suppresses ingestive behaviour [41,42]. We are only aware of two reports relating urocortin 3 to young-directed parental care. One is the poison frog study mentioned earlier [30] where urocortin 3 was upregulated in the preotic area of frogs performing parental care relative to non-parental ones. The other one revealed that urocortin 3 expressing neurons in the perifornical area of the hypothalamus are activated specifically during infant-directed aggression in both male and female mice and that optogenetic activation of these neurons triggers infant-directed aggression and neglect [43]. While aggression levels were not significantly different between our two treatments, they tended to be reduced after sensitization treatment. Furthermore, urocortin 3 expression did not covary with aggressive behaviour in our study. Rather than affecting infant-directed aggression in our system, urocortin 3 may serve different functions that support parental care, such as altered feeding patterns [44,45]. Interestingly, recent work proposes that molecules related to feeding behaviour may also be involved in complex social behaviours, such as parental care [46,47].

In the BnST only one gene was up-regulated in animals exposed to the sensitization procedure: APMAP. This was also the only gene up-regulated in the Tn if we consider a p_adj_ < 0.1 cut-off. The protein encoded by APMAP is important for white adipose tissue differentiation [48] and little is known regarding it’s neurobiological functions, except that is may be involved in suppressing amyloid-beta formation in brain [49]. The connection to parental care, if any, is unknown.

As no significant differences in gene expression was found between treatments in the Tn and no genes in this region showed covariation with brooding behaviour, this region is likely less critically important during the onset or production of brooding behaviour in birds.

Most of the differentially expressed genes that were down-regulated in the hypothalamus and BnST in animals that underwent the sensitization procedure, as well as those genes that negatively covaried with the expression of brooding behaviour, are considered immediate early genes [50] and are involved in regulation of transcription (FOSB, BTG2, FOS, NR4A3, TOB1, NPAS4, EGR1, EGR4, KLF2, JUN, SIK1) and in response to hormones (FOSB, FOS, NR4A3, NPAS4, JUN, SIK1). Very similar changes in immediate early gene expression patterns were found in several brain regions (including the hypothalamus; the BnST was not included in that study) in female mice, whereby immediate early gene expression was decreased in females during pregnancy and post-partum relative to virgin females [51]. One of the top enriched pathways highlighted through enrichment analysis was for response to corticosterone. Corticosterone is one of the main hormones involved in the response to stress in birds [52]. Data from a separate study on these birds showed that there were no differences between treatments in terms of serum corticosterone levels after the birds had been exposed to the novel chicks [12]. One possibility is that the way in which the hypothalamus and BnST respond to corticosterone changes in parental animals. Prolactin, for instance, has been implicated in the modulation of the stress response [53]. Reduced stress responsiveness may facilitate the acceptance of hatchlings in natural settings, which may otherwise be perceived as intruders. Alternatively, it is possible that the pathways highlighted reflect altered responsiveness to other hormones or peptides. For example, while central injection of neurotensin can increase c-Fos immunoreactivity in several brain nuclei (including hypothalamic nuclei) in non-parental rats [54], two studies that compared immediate early gene responses to neurotensin administration in maternal versus virgin female mice found a general decrease in reactivity (decreased Egr-1 and c-Fos production) in maternal females [55,56].

A previous study in female Japanese quail found that sensitized maternal females had increased c-Fos immunoreactivity in the BnST relative to sensitized non-maternal females [16]. This appears to be in contrast with our finding of higher FOS expression in the control relative to the sensitization treatment in this brain region. This difference may be related to the time point of parental care being studied and details of each study. In their study, the sensitization treatment consisted of exposure to chicks for 20 min periods over 5 consecutive days. The study of immediate early gene immunoreactivity was only done after the 5^th^ day. Also, on the 5^th^ day, after being allowed to interact with the chicks for 20 min, females were separated from the test chicks for 1 h prior to brain collection. These two major differences alone could result in different outcomes in our studies. The [16] study may be quantifying parental behaviour after is has been established, while our study reflects the onset of parental behaviour. Additionally, as c-Fos protein can be detected at 20 to 90 min after the stimulus [57], the separation from the chicks for 1 h prior to sample collection could have reflected responses related to maternal separation rather than maternal behaviour.

Our study indicates that both sexes possess the neural circuitry needed to produce brooding behaviour and that the changes in gene expression associated with this behaviour do not differ per sex. It is, however, important to note that our study was done under a background of low levels of gonadal hormones (short photoperiod), which may make males and females less different. It will therefore be interesting in future studies to assess how the photoperiod affects genomic responses associated with parental care behaviour and whether it introduces differences between the sexes.

## 5 Conclusion

Recent studies have used transcriptomic approaches to compare gene expression in specific brain areas as female birds transition from territorial defence to egg incubation [58], or from egg-laying to incubation behaviour [59]. Still, the range of neural mechanisms involved in post-hatching care in birds is virtually unknown. Our findings revealed new potential modulators of avian young-directed parental care, which deserve to be examined through further mechanistic studies and explored in other bird species and other taxonomic groups. Particularly exciting are neurotensin and urocortin 3, which potentially show conserved roles in parental care in mammals, amphibians, and birds, but are extremely under-explored in the context of parental behaviour. Natural variation in avian parental behaviour in response to young extends from abandoning the nest to alloparenting or even adoption of non-offspring deposited by brood parasites. The genes found to be associated with the onset of parental care and with variation in young-directed care in our study serve as a foundation for future studies aimed at investigating the neural underpinnings of variation in parental care within and across species.

## Ethics

Animal use and experimental design were approved by the Chapman University Institutional Animal Care and Use Committee (protocol # 2019-01).

## Data accessibility

RNA-seq mapping statistics, lists of DEGs and lists of enrichment terms have been uploaded as part of the supplementary material. Sequencing datasets generated and analysed during the current study are deposited in the NCBI Gene Expression Omnibus (GEO) repository, with record GSE162378 [dataset will be made publicly available once manuscript is accepted for publication].

## Authors’ contributions

PCL conceived the study, helped RDB carry out the animal experiments, collected the brain samples, performed the molecular work, the data analysis and wrote the manuscript. RDB carried out the animal experiments. All authors read and edited the manuscript.

## Competing interests

We declare we have no competing interests.

## Funding

This work was supported by start-up funding from Chapman University to PCL; RDB was supported by a Chapman University Grand Challenges Initiative fellowship.

## Acknowledgements

We thank Kaelyn Bridgette for her help with animal husbandry and for NEST lab undergraduate students for support in sample processing and behavioural analysis. We thank Greg Goldsmith for providing spontaneous parental care, particularly during the pandemic.

## Notes

### Competing Interest Statement

The authors have declared no competing interest.

### Summary of Updates

Both main text and supplementary files were updated.

